# Genomic history and forensic characteristics of Sherpa highlanders on the Tibetan Plateau inferred from high-resolution genome-wide InDels and SNPs

**DOI:** 10.1101/2021.06.23.449553

**Authors:** Mengge Wang, Weian Du, Renkuan Tang, Yan Liu, Xing Zou, Didi Yuan, Zheng Wang, Jing Liu, Jianxin Guo, Xiaomin Yang, Jing Chen, Meiqing Yang, Xianpeng Zhang, Lan-Hai Wei, Haibing Yuan, Hui-Yuan Yeh, Chuan-Chao Wang, Chao Liu, Guanglin He

## Abstract

Sherpa people, one of the high-altitude hypoxic adaptive populations, mainly reside in Nepal and the southern Tibet Autonomous Region. The genetic origin and detailed evolutionary profiles of Sherpas remain to be further explored and comprehensively characterized. Here we analyzed the newly-generated InDel genotype data from 628 Dingjie Sherpa people by merging with 4222 worldwide InDel profiles and collected genome-wide SNP data (approximately 600K SNPs) from 3324 individuals in 382 modern and ancient populations to explore and reconstruct the fine-scale genetic structure of Sherpas and their relationships with nearby modern and ancient East Asians based on the shared alleles and haplotypes. The forensic parameters of 57 autosomal InDels (A-InDels) included in our used new-generation InDel amplification system showed that this updated InDel panel is informative and polymorphic in Sherpas, suggesting that it can be used as the supplementary tool for forensic personal identification and parentage testing in the highland East Asians. Descriptive findings from the PCA, ADMIXTURE and TreeMix-based phylogeny suggested that Sherpas showed excess allele sharing with neighboring Tibeto-Burman Tibetans. Furthermore, patterns of allele sharing in *f*-statistics demonstrated that Sherpa people had a different evolutionary history compared with their neighbors from Nepal (Newar and Gurung) but showed genetic similarity with 2700-year-old Chokhopani and modern Tibet Tibetans. *QpAdm/qpGraph*-based admixture sources and models further showed that Sherpa, core Tibetans and Chokhopani formed one clade which could be fitted as having the main ancestry from late Neolithic Qijia millet farmers and other deep ancestries from early Asians. Chromosome painting profiles and shared IBD fragments inferred from FineStructure and ChromoPainter not only confirmed the abovementioned genomic affinity patterns but also revealed the fine-scale microstructures among Sino-Tibetan speakers. Finally, natural-selection signals revealed via iHS, nSL, and iHH12 showed signatures associated with disease susceptibility in Sherpa people. Generally, we provided the comprehensive landscape of admixture and evolutionary history of Sherpa people based on the shared alleles and haplotypes from the low-density forensic markers and high-density genome-wide SNP data. The more detailed genetic landscape of Sherpa people should be further confirmed and characterized via ancient genomes or single-molecule real-time sequencing technology.

## 1. Introduction

Sherpa is one of the Tibeto-Burman-speaking groups native to the most mountainous regions of China, Nepal, India, and Bhutan. Sherpa highlanders are renowned as excellent mountaineers and guides for their hardiness, expertise, and experience at high altitudes, who are considered as a good candidate for illuminating the genomic mechanism of high-altitude adaptation and have played an invaluable role in the exploration of peopling processes of the Himalayas [1–3]. However, when and where the Sherpa people originated and settled in the high-altitude regions of the Himalayas remains unclear. Besides, their genetic relationships with adjoining Tibetans also keep contentious and more genomic evidence was needed to support or disprove the hypothesis of the common origin of Sherpas and Tibetans. Genetic observations from matrilineal lineages demonstrated that Tibetans or Tibetan-related ancestors contributed considerably to the gene pool of Sherpa people [2, 4, 5]. A previous study of mitochondrial DNA (mtDNA) diversity on Sherpa reported considerable South Asian genetic components existing in Zhangmu Sherpas [6], but maternal lineages of other Sherpas living in Nepal and Tibet Autonomous Region of China revealed that matrilineal haplogroups with South Asian origin occurred in these studied Sherpas with a minor frequency [4]. Genetic findings based on Y-chromosomal single nucleotide polymorphisms (SNPs) showed that Sherpas shared most of their paternal lineages with indigenous Tibetans, as haplogroups D and O were their dominant founding paternal lineages [4]. Among non-uniparental markers, the genetic structure of the Nepal Sherpa and neighboring Nepalese populations based on the genome-wide SNPs indicated that the Sherpas were a remarkably isolated population, with little gene flow from surrounding Nepalese populations [7]. Further well-characterized genomic landscape of Tibeto-Burman and Indo-Aryan communities from the remote Nepalese valleys also provided evidence for Sherpa isolation [8]. A recent study focused on Tibetan highlanders in China based on the whole-genome sequencing data indicated that a small portion of Tibetans’ and Sherpas’ ancestral components originated from separate ancient populations, which were estimated to have lived approximately 7000 to 11000 years before present (YBP) [9]. Lu et al. also demonstrated that Sherpas and Tibetans shared more recent ancestors than one of them with Han Chinese [9]. Conversely, Jeong et al. pointed out that Sherpa and Han Chinese served as dual ancestral populations of Tibetans and supported that modern Tibetans were the recent admixture result of ancestral sources related to Sherpas and Hans [10]. Generally, some genetic evidence supported the Nepal-Tibet migrations, while some others supported the Tibet-Nepal migrations; some genetic findings suggested that Tibetans were the ancestral populations of the Sherpas, while some others indicated that Tibetans were a mixture of ancestral populations related to the Sherpa and Han Chinese. The disparity in these perspectives reflects a fundamental division in the demographic history of Sherpas, suggesting that the genetic makeup of Sherpas remains to be elucidated.

Notably, genetic researches focused on the Sherpas were mainly performed in the fields of archaeology [11, 12], molecular anthropology [7–9], medical genetics [3, 5], and genetic genealogy [2, 4], only a few forensic-related studies focusing on the Nepal Sherpas were conducted based on the low-density short tandem repeats (STRs) [13, 14]. Insertion and deletion polymorphisms (InDels), the second most abundant polymorphism across the human genome with low mutation rates and small amplicon lengths [15], combine the desirable features of both SNPs and STRs. The prominent properties enable this molecule to be genotyped via capillary electrophoresis (CE) platform. In addition, InDels cause similar levels of variation as SNPs and do not involve repetitive sequences, which can avoid stutter artifacts that may complicate the interpretation of STR profiles [16]. Reportedly, the 1000 Genomes Project has characterized ~3.6 million short InDels [17] and the analysis of high-coverage genome sequences of 54 diverse human populations included in the HGDP-CEPH panel identified ~8.8 million small InDels [18], which promoted the practical application of InDel loci in forensic investigations [19–21]. In the last few years, several commercialized InDel kits have been designed, such as the Investigator DIPplex kit developed by Qiagen [22–24], the AGCU InDel 50 kit [20, 25, 26], and the AGCU InDel 60 kit developed by AGCU ScienTech Incorporation. However, the practicability and efficiency of the updated AGCU InDel 60 kit have not been validated in geographically/ethnically/linguistically different populations.

Here, we first sought to explore the forensic characteristics and genetic relationships of the Sherpa people and their neighbors on the Tibetan Plateau using the updated AGCU InDel 60 kit. Next, we comprehensively characterized the evolutionary genomic history of Sherpas based on one of the most representative modern and ancient genome-wide datasets from two analysis strategies: allele frequency-based shared alleles (PCA, ADMIXTURE, and *f*-statistics) and haplotype-based shared ancestry chunks (ChromoPainter and FineStructure). The sampled Sherpas live in Dingjie County, Shigatse, Tibet Autonomous Region, which borders Sankhuwasabha and Taplejung Districts of Nepal to the south and the Sikkim State of India to the southeast and is one of the four counties that comprise the Qomolangma National Nature Preserve. We demonstrated forensic effectiveness of our used InDel panel in highland adaptive Sherpa and found a strong genetic affinity between Sherpas and Tibetans, all of them primarily originated from Neolithic Yellow River farmers.

## 2. Materials and methods

### 2.1. Sample preparation, DNA extraction, and genotyping

Bloodstain samples from 250 males and 378 females were collected from Sherpa individuals living in Dingjie County, Shigatse, Tibet Autonomous Region after receiving written informed consent. All participants were required to be the indigenous Sherpas whose ancestors have lived in Tibet Autonomous Region for at least three generations. This study was approved by the Ethical Committee of North Sichuan Medical College and Xiamen University, and all procedures were performed following the recommendations of the Declaration of Helsinki [27].

Human genomic DNA (gDNA) was extracted using the QIAamp DNA Mini Kit (QIAGEN, Germany) and quantified using the Qubit dsDNA HS Assay Kit (Thermo Fisher Scientific) on an Invitrogen Qubit 3.0 fluorometer following the manufacturer’s recommendations. The gDNA was subsequently diluted to 2.0 ng/μl and stored at −20°C until amplification. The control DNA 9947A and 9948 were adopted as reference samples.

Fifty-seven A-InDels and three sex-determination loci (rs76041101, rs199815934, and Amel) were amplified simultaneously using the AGCU InDel 60 kit on a ProFlex 96-Well PCR System (Thermo Fisher Scientific) following the manufacturer’s protocol. The rough physical positions of all genotyped loci are provided in **Figure 1A**. The Applied Biosystems 3500xl Genetic Analyzer was utilized to separate the amplified products, and the GeneMapper ID-X v.1.5 software (Thermo Fisher Scientific) was applied to allocate alleles.

**Figure 1.**
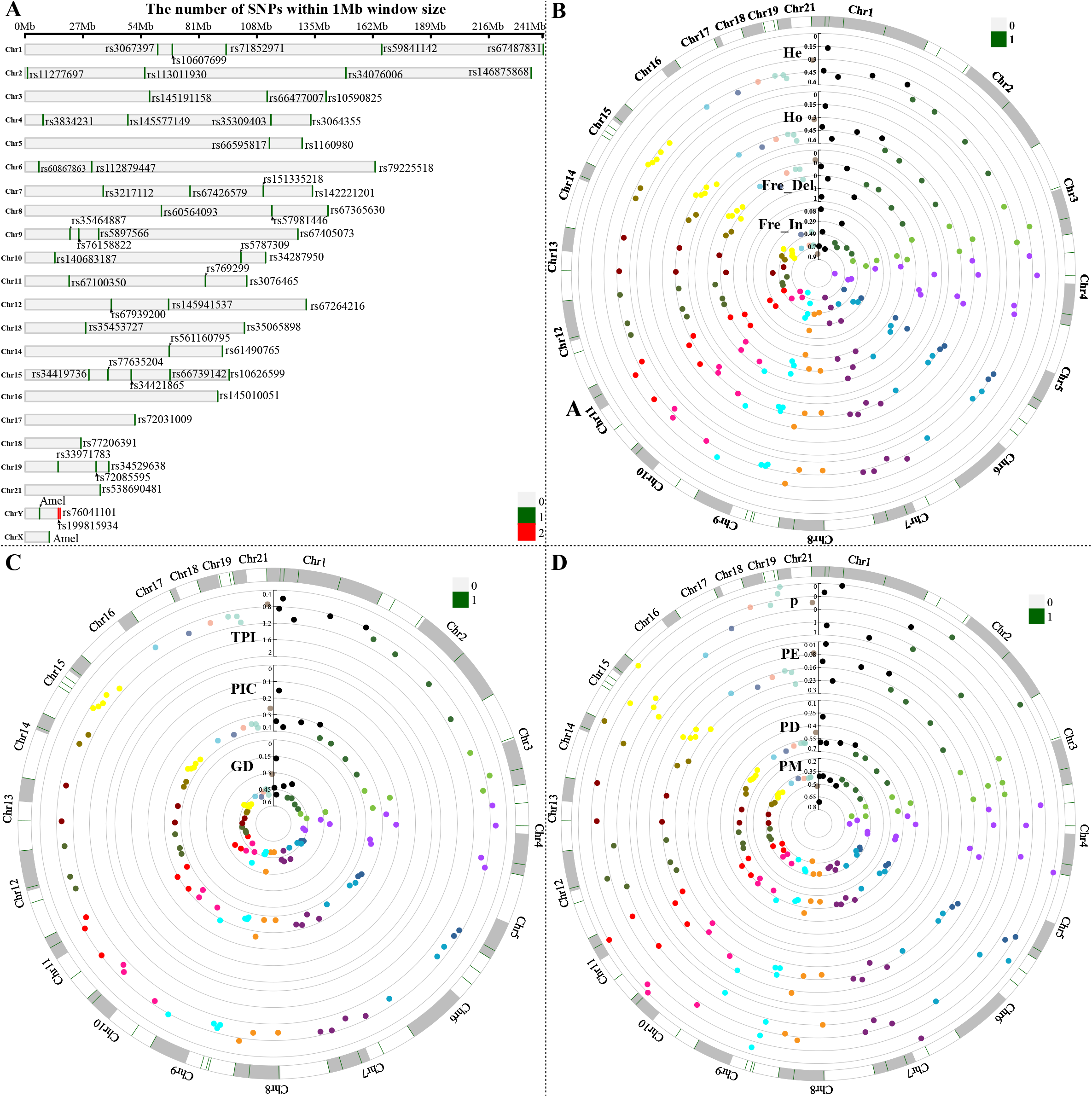
Our used InDel markers and their forensic statistical parameters. (A) Distributions of 60 InDels in the human chromosomes. (B-D) Allele frequency distribution and forensic statistical parameters of 57 autosomal InDels in the Sherpa people.

### 2.2. InDels and genome-wide SNP dataset composition

To reconstruct the genomic history of Sherpas and investigate the genetic relatedness between the studied Sherpas and reference populations, the newly generated data were merged with genotypes of 57 A-InDels retrieved from the 1000 Genomes Phase III release [17]. Furthermore, genotype data of 39 overlapped A-InDels from four Sinitic-(Han and Hui), five Tibeto-Burman-(Tibetan, Yi, and Mosuo), one Turkic-(Uyghur), and two Tai-Kadai-speaking (Gelao and Li) populations [26, 28] were merged into the above-mentioned 57-A-InDel-based dataset. The detailed information of reference populations is listed in **Table S1** and the geographical locations of all involved populations are displayed in **Figure 2A**. We then collected genome-wide SNPs from Eurasian modern populations genotyped via Human Origins array [29–31] and collected eastern Eurasian ancient population data as the direct ancient ancestral sources to explore the genetic contribution and possible existing genetic continuity and admixture process. Ancient Chinese populations from Nepal, Mongolian Plateau, the Yellow River Basin and the Yangtze River Basin were our main focuses [32–36].

**Figure 2.**
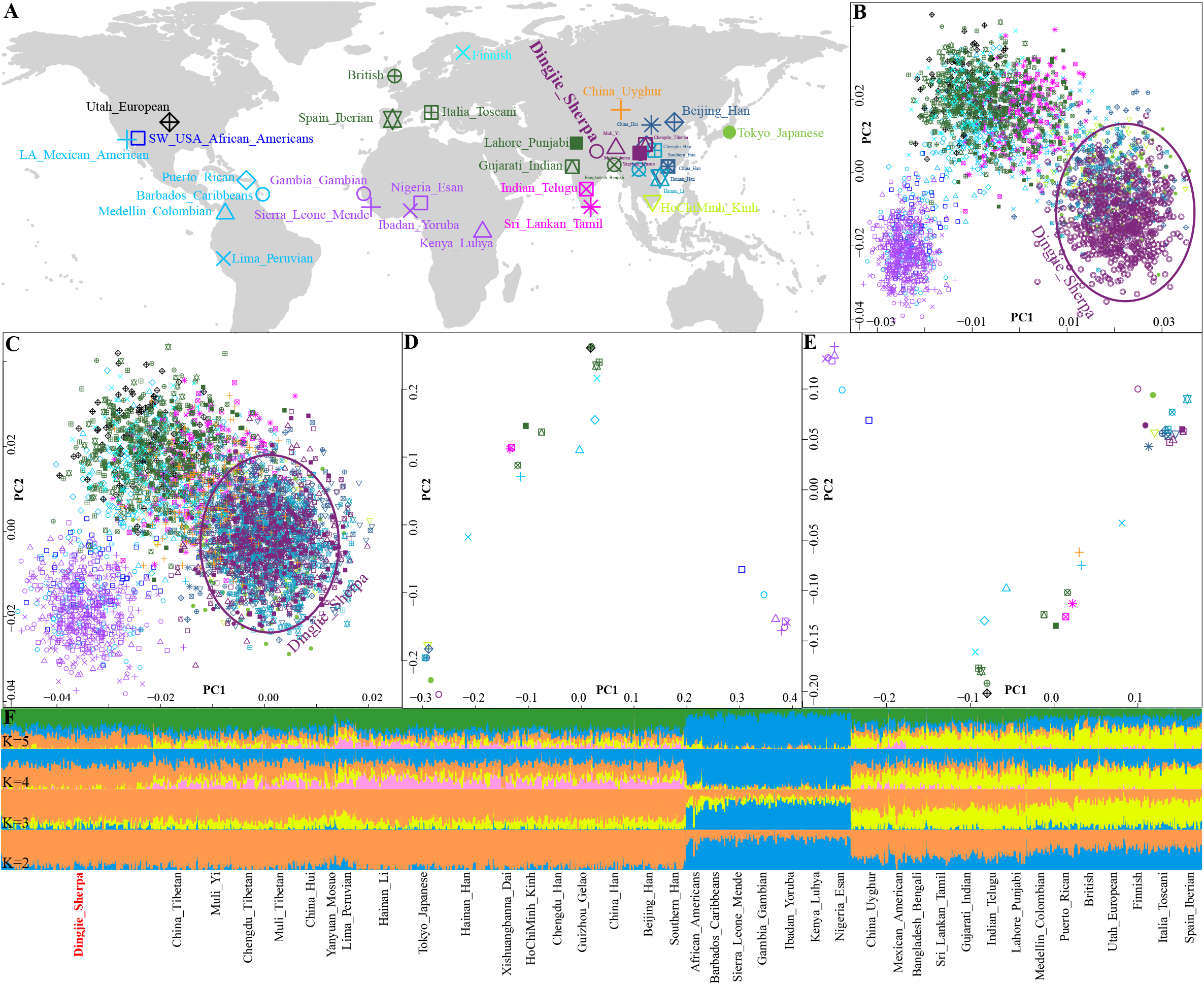
Geographical information and genetic structure based on InDel markers. (A) The geographical positions of the studied Sherpa population in Dingjie country, southern Tibet Autonomous Region. (B) Principal component analysis (PCA) among 27 worldwide populations based on the top three components extracted from the genetic variations of 57 InDels. (C) PCA results among 4850 individuals from 39 worldwide populations based on the genetic polymorphisms of 47 InDels. (D-E) PCA analysis results among worldwide populations based on the allele frequency distribution based on two InDel datasets. (F) Model-based ADMIXTURE results showed the genetic composition of 27 worldwide populations with the K values ranging from 2 to 5. The model with the predefined five ancestral sources had the less cross-validation errors.

### 2.3. Data analysis

#### 2.3.1. InDel-based analysis of genetic variations

The exact tests of linkage disequilibrium (LD) and Hardy-Weinberg equilibrium (HWE), as well as the estimations of observed heterozygosity (Ho) and expected heterozygosity (He) were conducted using the Arlequin v.3.5.2.2 [37]. The browser-based software of STR Analysis for Forensics (STRAF) was adopted to calculate the allele frequencies and forensic statistical parameters: discrimination power (DP), match probability (MP), probability of exclusion (PE), polymorphism information content (PIC), and typical paternity index(TPI) [38]. We used the Gendist package built into the PHYLIP v.3.698 to estimate Nei’s genetic distances [39]. We subsequently conducted genotype-based principal component analysis (PCA) using the genome-wide Complex Trait Analysis (GCTA) tool [40] and frequency-based PCA using the Multivariate Statistical Package (MVSP) v.3.22 [41]. The model-based clustering analysis was performed via the ADMIXTURE v.1.3.0 [42] with 10-fold cross-validation (cv) in 100 bootstraps to dissect the ancestral components of target Sherpas. To infer the patterns of population splits and mixtures in multiple populations, we ran TreeMix v.1.13 [43] with migration events ranging from 0 to 10 to build a maximum likelihood tree. We also conducted the analysis of molecular variance (AMOVA) using the Arlequin v3.5.2.2 by grouping involved populations according to geography, language family, or ethnicity classifications.

#### 2.3.2. Allele-based sharing ancestry inferred from the high-density genome-wide data

Focused on the high-density SNP dataset, we used the smartpca program built into the EIGENSOFT v.6.1.4 [29] to carry out PCA based on the Chinese modern populations and their genetically homogeneous neighbors, and ancient individuals were projected onto the top two components. PLINK v.1.9 [44] was applied to prune SNPs in strong linkage disequilibrium with the following parameters (--indep-pairwise 200 25 0.4). We then conducted model-based clustering analysis using the ADMIXTURE v.1.3.0 [42] with 10-fold cross-validation (--cv = 10) and predefined ancestry sources ranging from 2 to 20. We also built a bifurcating tree with the maximum likelihood algorithm based on the genome-wide allele frequency data using the TreeMix [45].

We used the *qp3Pop* program [29] of the ADMIXTOOLS to perform outgroup*-f_3_*(Source, Sherpa; Mbuti) to explore the shared genetic drift and conduct admixture*-f_3_*(Source1, Source2; Sherpa) to explore the status of genetic admixture and corresponding proxy ancestral sources. We further used the *qpDstat* package [29] of the ADMIXTOOLS to perform *f_4_*-statistics in the forms *f_4_*(Eurasian1, Eurasian2; Sherpa, Outgroup) and *f_4_*(Eurasian1, Sherpa; Eurasian2, Mbuti) to investigate genetic affinity, admixture, and continuity with different comparative populations. The potential ancestral sources and corresponding admixture proportions were estimated using the *qpWave/qpAdm* packages [29] and graph-based admixture modeling was conducted using the *qpGraph* software [29].

#### 2.3.3. Shared haplotype chunks inferred from the FineStructure and ChromoPainter

Haplotype phasing was carried out by means of the statistical-based method implemented in the SHAPEIT [46] using default parameters. The phased haplotype data were used to paint the target individual’s chromosome using all other included genetic material from other populations as the potential donor populations. The inference framework of chromosome painting of recipient chromosomes in the context of extensive donor chromosomes was used and conducted via a statistical algorithm instrumented in the ChromoPainter and ChromoCombine [47]. Individual-level model-based Bayesian clustering was carried out using the FineStructure v.4 [47] based on the output of ChromoPainter. The model-based likelihood and PCA based on our used coancestry matrix were estimated using the R packages. Finally, natural selection signals were explored via selscan based on the phased haplotype data and Identity by Descent (IBD) fragments were calculated using fineIBD [46].

## 3. Results

### 3.1. Overview of forensic characteristics and population genetic structure inferred from the updated InDel panel

#### 3.1.1. Allele frequency distribution and forensic statistical parameters

The present study has generated the first batch of InDel-based genotype data of 628 Sherpa individuals (**Table S2**). The allele frequency and corresponding forensic parameters are listed in **Table S3** and plotted in **Figures 1B-1D**. Two InDel loci (rs3064355 and rs5787309) showed significant departure from HWE after conducting Bonferroni correction (**Table S3**, p < 0.0009), five pairs of InDels (rs5787309 and rs34287950, rs67426579 and rs151335218, rs5897566 and rs76158822, rs35464887 and rs76158822, rs72085595 and rs34529638) located on the same chromosome showed LD after applying Bonferroni correction (**Table S4**, p < 0.00003). The allele frequencies of insertions fluctuated within 0.1449-0.9076. The values of Ho and He ranged from 0.1656 (rs10607699) to 0.5526 (rs34529638) and 0.1678 (rs10607699) to 0.5004 (rs146875868). The values of DP and PE varied from 0.2921 (rs10607699) to 0.6319 (rs145941537) and 0.0211 (rs10607699) to 0.2378 (rs34529638). The total probability of discrimination power (TDP) value reached 1-1.5732E-22, and the combined probability of exclusion (CPE) value was approximately 0.9999. The measured value of PIC was in the range of 0.1536 (rs10607699) to 0.3750 (rs146875868), the minimum value of TPI was 0.5992 at rs10607699, whereas the maximum reached 1.1174 at rs34529638.

#### 3.1.2. Genetic similarity and differences between studied Sherpas and reference populations

Genetic affinities between studied Sherpas and reference populations were explored via Nei’s genetic distances, PCA, ADMIXTURE, and TreeMix analyses. The 57-InDel-based genetic distances indicated that studied Sherpa people were genetically close to East Asian reference populations (**Table S5**), including northern and southern Han Chinese, Yunnan Dais and Japanese. The 39-InDel-based genetic distances revealed that studied Sherpas were genetically close to previously published Tibeto-Burman-speaking populations residing in the Tibetan Plateau (**Table S6**, Yanyuan_Mosuo: 0.0248, Muli_Tibetan: 0.0269, Muli_Yi: 0.0298). The 57-InDel-genotype-based and 39-InDel-genotype-based PCAs (**Figures 2B-2C** and **S1A-S1B**) only identified three rough clusters: African cluster, East Asian cluster, and a mixed cluster including European, American, and South Asian populations. Generally, the studied Sherpas were overlapped with Sino-Tibetan-speaking individuals. The frequency-based PCAs (**Figures 2D-2E** and **S1C-S1D**) identified four geography-related clusters: African cluster, European cluster, South Asian cluster, and East Asian cluster, the American populations were scattered along the PC2 or PC3. Specifically, the studied Dingjie Sherpa showed a close genetic relationship with East Asian reference populations, especially Sino-Tibetan-speaking populations and Tokyo Japanese.

We further performed model-based ADMIXTURE analysis to roughly model the ancestral components of the target Dingjie Sherpa (**Figure 2F**). At K = 2, we observed two distinct ancestral components deriving from non-African (orange component) and African populations (blue component). At K = 3, the yellow ancestral component occurred in Chinese Uyghur, South Asian, American, and European populations. As the K values increased, two additional ancestries, including East Asian-dominant ancestral component (showed in green) and non-Dingjie_Sherpa-related ancestral component (showed in pink), were observed. Obviously, ancestral compositions of the studied Sherpas were similar to other Tibeto-Burman speakers. Furthermore, we built maximum likelihood topologies to model the gene flow between target populations and proxy sources. The 39-InDel-based phylogenetic tree demonstrated that the Dingjie Sherpa grouped with Tibeto-Burman-speaking populations and showed strong genetic affinities with Tibetans (**Figures 3A-3B**). The 57-InDel-based topology revealed that the Dingjie Sherpa clustered with East Asian reference populations and showed a relatively close genetic relationship with Tokyo Japanese (**Figures 3C-3D**). Moreover, we observed gene flow events from Gambia_Gambian-related ancestry into East Asians, from Lima_Peruvian-related ancestry into Medellin_Colombian, from European-related ancestry into Mexican_American, and from Utah_European-related ancestry into Lahore_Punjabi. To explore the factors that might have played roles in shaping the InDel diversity among geographically, linguistically and ethnically diverse populations, we conducted a series of AMOVA analyses and found that the among-group variations were slightly higher when dividing worldwide populations into geographically different groups, compared with that among linguistically different groups (**Table S7**). We also observed that the among-group variations were slightly higher when dividing East Asian populations into ethnically different groups, compared with that among language-related and altitude-related groups.

**Figure 3.**
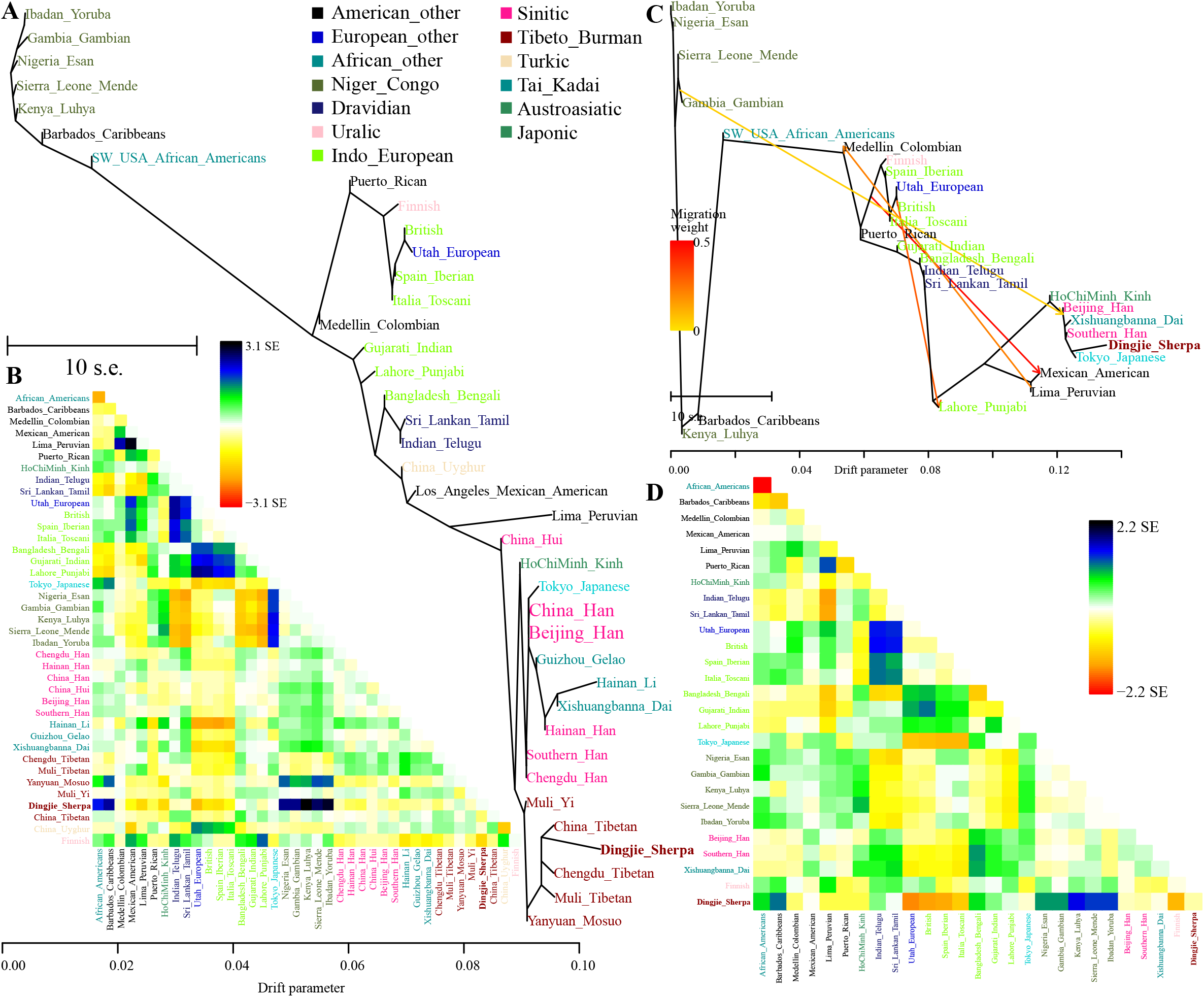
Phylogenetic relationship and residual values among worldwide populations. (A-B) TreeMix-based phylogenetic relationship and residual values among 39 populations with zero migration event. (C-D) TreeMix-based phylogenetic relationship and residual values among 27 populations with four migration events. Populations were color-coded based on their geographical affinity and linguistic affinity.

### 3.2. Genetic origins of Sherpa people from the Tibetan Plateau inferred from genome-wide SNP data

#### 3.2.1. Genetic structure and affinity inferred from PCA, ADMIXTURE, and TreeMix-based phylogeny

Although the forensic used genetic markers, such as STRs and InDels, possessed powerful statistical performance in the personal identification and parentage testing, the limited resolution (the number of the genetic markers is low) of deep ancestry dissection has hindered its usefulness in molecular anthropology and population genomic history reconstruction. Thus, to gain new insight into the genetic makeup of Sherpa highlanders, we retrieved genome-wide SNP data of Sherpa people from one of the most comprehensively representative datasets, including all publicly available ancient and modern Eurasian genomes [30, 31, 34, 36, 48]. We explored the genetic admixture history based on the sharing independent alleles and shared haplotype chunks consisting of the successive SNPs. We also provided the direct genetic relationship between Sherpa people and their geographically close ancient ancestors from the Tibetan Plateau (Chokhopani, Samdzong and Mebrak [33]), Yellow River millet farmers (Qijia Jinchankou and Lajia people [36], Henan Yangshao and Longshan people [36], and Shandong Houli people [35]) and other ancient East Asians [34].

To explore the general patterns of genetic relatedness between the target Sherpas and reference East Asian populations, we first performed PCA analyses based on all East Asian populations and all Sino-Tibetan sub-populations. All published Sino-Tibetan (Tibeto-Burman and Sinitic) people were our focus to characterize the genetic history and relationship of Sherpas and their neighbors (**Figure 4A**). The HO-based East Asian PCA (**Figure 4B**) identified two genetic clines: the southern cline consisting of Austroasiatic-, Tai-Kadai-, Austronesian-, Tibeto-Burman-(mainly in Southeast Asia), and Hmong-Mien-speaking populations, and the northern one consisting of the studied Sherpas, Sinitic-, Japonic-, Koreanic-, Tibeto-Burman-(mainly in the Tibetan Plateau), Tungusic-, Mongolic-speaking populations, and ancient individuals in Russia and Mongolia. Tibeto-Burman people from the Tibetan Plateau and mainland Southeast Asia were separated into two clusters (purple color-coded). We observed that the target Sherpas clustered with Tibetan populations. The Sino-Tibetan-related PCAs (only including 361 people from 44 Sino-Tibetan-speaking populations) based on top three components revealed clear population subclusters within 44 Sino-Tibetan-speaking populations: Newar-related, Karen-related, low-altitude Sino-Tibetan-related, and high-altitude Tibeto-Burman-related genetic clines (**Figures 4C-4E**). The Sherpa people showed close genetic relationships with Tibet Tibetans (especially Shigatse Tibetan and Shannan Tibetan).

**Figure 4.**
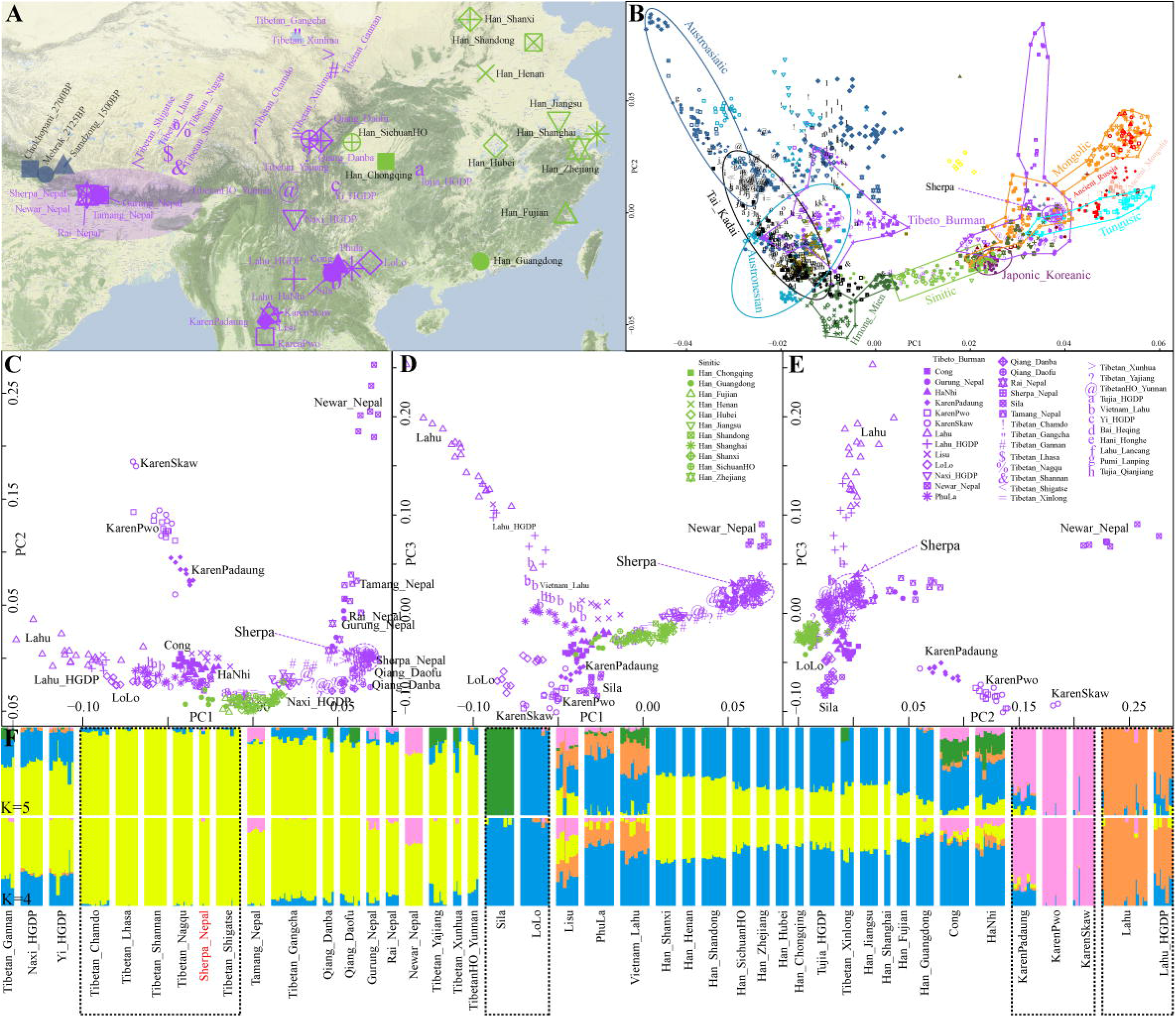
Genetic structure and geographical position of populations used in the population genomic analysis based on the genome-wide data. (A) Geographical positions of 44 Sino-Tibetan populations and three Nepal ancient populations, five Nepal populations (including one Sherpa population) were emphasized. (B) PCA analysis among 1612 modern and ancient East Asians based on the Human Origins, ancient populations from Nepal, China, Japan, Mongolia and southern Siberia were projected onto the top two components. (C-E) PCA results among 361 Sino-Tibetan-speaking individuals from 44 populations. (F) Ancestry composition of 361 Sino-Tibetan people based on unlinked SNPs.

Clustering patterns inferred from the ADMIXTURE results among Sino-Tibetan-speaking populations revealed multiple sources of their ancestral components (**Figure 4F**). Under the model of four predefined ancestral sources with the less cross-validation error (K = 4), we found four specific ancestral components that were respectively maximized in Tibet Tibetans and target Sherpas (yellow), Sila and LoLo (blue), Karen-related populations (pink), and Southeast Asian Lahu-related populations (orange). Model-based clustering of five predefined ancestral sources confirmed the Sila-specific ancestry, which also occurred in relatively low-altitude Tibetan populations (except Yunnan Tibetan), Qiang people, Lisu, PhuLa, Vietnam Lahu, Cong, and HaNhi. Model-based admixture results showed a similar genetic composition between Sherpas and Tibetans, suggesting their close genetic relationship and shared common ancestral evolutionary history. The maximum likelihood trees with migration events indicated that the target Nepal Sherpa was aggregated with high-altitude Tibetan populations to form the high-altitude-related clade, which then clustered with Qiang and relatively low-altitude Tibetan populations (**Figure S2**), which further confirmed the similar gene pool of Sherpas and Tibetans. Here, we also identified evidence of the documented gene flow from Newar to Southeast Asian Tibeto-Burman-speaking populations, from Lahu to the common ancestor of LoLo, PhuLa and Lahu, from Tamang to the common ancestor of Sherpas and central and southern Tibet Tibetans (Lhasa Tibetan, Shannan Tibetan, and Shigatse Tibetan), from Guangdong Han to Fujian Han, which suggested that extensive genetic admixture occurred among Sino-Tibetan populations shaped these genetically attested diversities.

#### 3.2.2. Shared alleles inferred from f-statistics

To explore the inter-population genetic relatedness between the studied Nepal Sherpa and other East Asians, we estimated pairwise Fst genetic distances among 44 Sino-Tibetan-speaking groups (**Table S8**). We found that the Nepal Sherpa had close genetic relationships with Tibetans on the Tibetan Plateau (Shannan Tibetan: 0.0036, Lhasa Tibetan: 0.0040, Shigatse Tibetan: 0.0043, Nagqu Tibetan: 0.0047, and Chamdo Tibetan: 0.0059). We subsequently measured the shared genetic drift (**Table S9**) via the outgroup *f_3_*-statistics of the form *f_3_*(Target population of Sherpa, Reference modern and ancient populations; Mbuti), in which large *f_3_* values denoted more shared ancestry and a close genetic relationship between target and reference populations. We observed that the Nepal Sherpa possessed the most shared alleles with Qiang groups (Qiang_Danba: 0.2894, Qiang_Daofu: 0.2886), high-altitude Tibetan populations (Tibetan_Lhasa: 0.2886, Tibetan_Nagqu: 0.2885, Tibetan_Shigatse: 0.2884, Tibetan_Shannan: 0.2883, and Tibetan_Chamdo: 0.2882), and middle/late Neolithic ancient populations (Wuzhuangguoliang_LN: 0.2902, Jinchankou_LN: 0.2888, and Miaozigou_MN: 0.2883). We also performed admixture *f_3_*-statistics of the form *f_3_*(Source1, Source2; Target populations) to identify plausible ancestral sources of the Nepal Sherpa (**Table S10**). Surprisingly, we did not observe significant negative Z-scores (< −3), which denoted that there was no evidence to support that Nepal Sherpa was descended from a recent genetic admixture event of our used modern and ancient source1 and source2. No-fitted ALDER-based curves and models have further confirmed the relatively isolated genetic structure of Sherpa people. We found that only the combination of (Mebrak_2125BP, BanChiang_IA) could produce relatively negative Z-scores (< −1.5), indicating their close genetic relationship.

Four population-based analyses provided strong power to validate our proposed topologies and potential existing gene flow events and gene flow directions [29]. Thus, we also conducted *qpWave* analyses to validate the genetic continuity between Nepal ancients and Sherpas, and Sherpas and Tibetans using one set of distinct outgroups (**Figure 5A**) and more powerful outgroups (**Figure 5B**). P values of pairwise qpWave large than 0.05 denoted that two left populations could be explained via one common ancestral source without additional gene flows compared with our used outgroups (distant outgroups: Mbuti, Ust_Ishim, Kostenki14, Papuan, Australian, Mixe, MA1, Onge, Atayal and Yamnaya_Samara; additional outgroups in more powerful outgroups: Mongolia_N_East, Bianbian_EN, Wuqi_EN, Qihe_EN, Liangdao2_EN, Yumin_EN and Xiaowu_MN). Genetic continuity was identified between Sherpa people and 2700-year-old Chokhopani and 2125-year-old Mebrak. Besides, genetic similarity between Sherpas and Shannan, Shigatse, Lhasa and Naqu Tibetans were also confirmed via pairwise qpWave analysis (**Figure 5B**). Additionally, we performed a series of symmetry-*f_4_*-statistics of the form *f_4_*(Eurasian1, Eurasian2; Target Nepal Sherpa, Mbuti) and found that the Nepal Sherpa shared more alleles with Tibetan groups, northern Han Chinese (Shanxi, Henan and Shandong), ancient populations (Samdzong_1500BP, Mebrak_2125BP, and Chokhopani_2700BP) on the Tibetan Plateau, and middle/late Neolithic ancient northern East Asians when compared with modern and ancient southern East Asians, Southeast Asians, South Asians, and ancient individuals not mentioned above, such as the most negative Z-score of *f_4_*(Sintashta, Tibetan_Shigatse; Sherpa_Nepal, Mbuti) = −62.023*SE. To clarify if additional ancestral sources contributed to the gene pool of Nepal Sherpa, affinity-*f_4_*-statistics of the form *f_4_*(Ancestral source candidate, Target Nepal Sherpa; Reference population, Mbuti) were conducted. When ancient individuals on the Tibetan Plateau and modern Tibetan groups were adopted as the ancestral sources, no significant negative Z-scores were observed in *f_4_*(High-altitude Tibetan populations/Mebrak_2125BP/Chokhopani_2700BP, Sherpa_Nepal; Reference population, Mbuti), which demonstrated that high-altitude Tibetans, Mebrak_2125BP, and Chokhopani_2700BP formed one clade with Sherpa people and these ancient populations might be the direct ancestors of the Nepal Sherpa (**Figure 5C**). Specifically, additional gene flow related to ancient Yellow millet farmers and modern northern East Asians into the Nepal Sherpa was identified in *f_4_*(Samdzong_1500BP, Sherpa_Nepal; Tibetan_Yajiang/HmongDaw/Han_Shandong/Mongolia_N_East/Dao/Mlabri/Tibetan_Gannan/Shimao_ LN, Mbuti), *f_4_*(Tibetan_Gangcha/Tibetan_Xunhua/Tibetan_Gannan, Sherpa_Nepal; Tamang_Nepal/Rai_Nepal/Gurung_Nepal, Mbuti), and *f_4_*(Tibetan_Gannan, Sherpa_Nepal; Newar_Nepal/Mongolia_N_East/Yumin_EN, Mbuti). The Z-scores of *f_4_*(Ancient East Asians, Sherpa_Nepal; Non-Tibetan-related reference population, Mbuti) revealed that the combinations of (Mongolia_N_East, Sherpa_Nepal; Tamang_Nepal/Rai_Nepal/Gurung_Nepal/Newar_Nepal/HtinPray, Mbuti) and (Yumin_EN, Sherpa_Nepal; Tamang_Nepal/Rai_Nepal/Atayal, Mbuti) could produce significant negative Z-scores.

**Figure 5.**
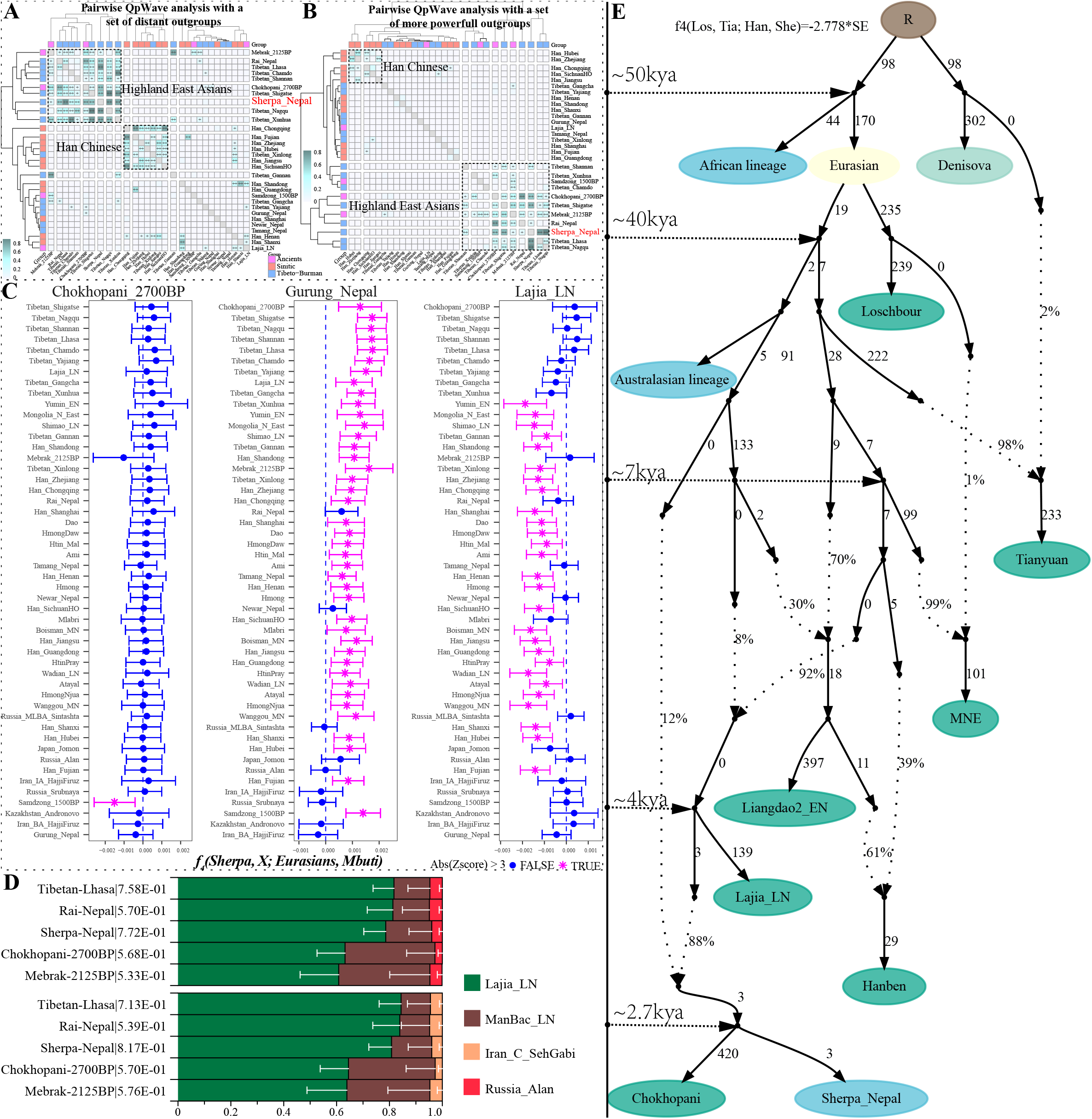
Population genetic analysis results based on the shared alleles in the combination of *f*-statistics. (A-B) Pairwise qpWave analysis based on distant outgroup set (Mbuti, Russia_Ust_Ishim, Russia_Kostenki14, Papuan, Australian, Mixe, Russia_MA1_HG, Onge, Atayal, Mongolia_N_East, Bianbian_EN, Wuqi_EN, Qihe_EN, Liangdao2_EN, Yumin_EN, Russia_EBA_Yamnaya_Samara and Xiaowu_MN) and more powerful outgroup set (Mbuti, Russia_Ust_Ishim, Russia_Kostenki14, Papuan, Australian, Mixe, Russia_MA1_HG, Onge, Atayal, Mongolia_N_East, Bianbian_EN, Wuqi_EN, Qihe_EN, Liangdao2_EN, Yumin_EN, Yamnaya_Samara_EBA and Xiaowu_MN) displayed genetic heterozygosity and homogeneity. (C) Four-population-based *f*-statistics in the form *f_4_*(Sherpa, Chokhopani_2700/Gurung Nepal/Lajia_LN; Eurasians, Mbuti) showed the genetic continuity or heterozygosity between Sherpas and their ancestors and modern neighbors. (D) QpAdm-based admixture results of modern and ancient highland East Asians used nine outgroups (Mbuti, Russia_Ust_Ishim_HG, Russia_Kostenki14, Papuan, Australian, Mixe, Russia_MA1_HG, Onge and Atayal). (E) Deep evolutionary history of modern Sherpas. Dotted line denoted the reconstructed admixture events and the corresponding admixture proportions were marked on it. The shared genetic drift of branch length was marked using 1000\**f_2_*. The time-scale was marked based on the recalibrated years of ancient samples, which was the simplified scale.

#### 3.2.3. Estimation of ancestral composition and admixture proportion

Considering the patterns of admixture events that were observed in the *f*-statistics results, we modeled the minimum number of proxy ancestral populations using the *qpWave* and estimated the corresponding ancestry proportions using the *qpAdm*. The two-way admixture models showed that the Nepal Sherpa could be modeled as an admixture of 0.246 ManBac_LN-like ancestry and 0.754 Lajia_LN-like ancestry (p_rank1: 0.0503), 0.543 ManBac_LN-like ancestry and 0.457 Mongolia_N_East-like ancestry (p_rank1: 0.0565), or 0.036 Sintashta-like ancestry and 0.964 Lajia_LN-like ancestry (p_rank1: 0.4077). Compared with the ancient populations on the Tibetan Plateau, the studied Nepal Sherpa possessed more Lajia_LN-like ancestral components. While no obvious differences in the proportion of Lajia_LN-like ancestry were observed between the Nepal Sherpa and high-altitude Tibetan groups. The results of three-way admixture models indicated that the Nepal Sherpa could be fitted as having 0.786 Lajia_LN-like ancestry, 0.174 ManBac_LN-like ancestry, and 0.040 Alan-like ancestry, or having 0.808 Lajia_LN-like ancestry, 0.152 ManBac_LN-like ancestry, and 0.040 Iran_C_SehGabi-like ancestry (**Figure 5D**). Similarly, we found that the studied Nepal Sherpa, high-altitude Tibetan groups, and Nepalese harbored more Lajia_LN-like ancestry than ancient groups on the Tibetan Plateau. We then used the *qpGraph* to further test if the phylogeny-based model with population splits and mixtures could provide a reasonable fit to the data, using the combination of all *f_4_*-statistics. We obtained the best-fitting model for the Nepal Sherpa with an absolute Z-score of 2.778 (**Figure 5E**). The optimal model revealed that the Nepal Sherpa had ~88% ancestry from the ancestor of Lajia_LN and ~12% ancestry from the Australasian lineage. Ancestral admixture sources and corresponding admixture proportions inferred from qpAdm and qpGraph consistently supported Sherpa people had a similar evolutionary history as Tibetan people, harboring the major ancestry of East Asians related to Neolithic millet farmers from the Yellow River Basin.

### 3.3. Finer-scale population demographic history and natural selection signals inferred from the shared haplotype

FineStructure model-based analyses based on the linked coancestry matrix of two different datasets were conducted to explore the population structure of Sherpas and their neighbors: one large dataset consisted of 361 people from 44 Sino-Tibetan populations and the other small one included 121 people from 18 Sinitic groups and highland Tibeto-Burman-speaking populations. Two-dimensional plots based on the top two components of haplotype-based PCA showed a clear separated position between Karen and other Sino-Tibetan people (**Figure 6A**). The third component separated Lahus from other included reference populations. The fourth and fifth components separated Sila, Lahu and Lolo populations from other people (**Figure 6B-J**). These separated populations were representative ancestral sources that harbored the maximized ancestry proportion in model-based ADMIXTURE. Sherpas were clustered together with Tibetans. Heatmap and the corresponding clustering patterns based on the coancestry matrix showed five major branches (**Figure 6K**): Han Chinese population branch was clustered closely with lowland Tibetan-Burman branch and then grouped with Lahu branch; Highland Tibetan-Burman speakers from the Tibet Autonomous Region and Nepal were separated. We also investigated interactions between populations by analyzing the number and length of shared IBD segments. The large and long shared IBD segments showed wide interaction and/or recent common ancestor sharing of the Nepal Sherpa with Tibetan groups, Tamang and Rai from Nepal, and Qiang groups (**Figure 6L**).

**Figure 6.**
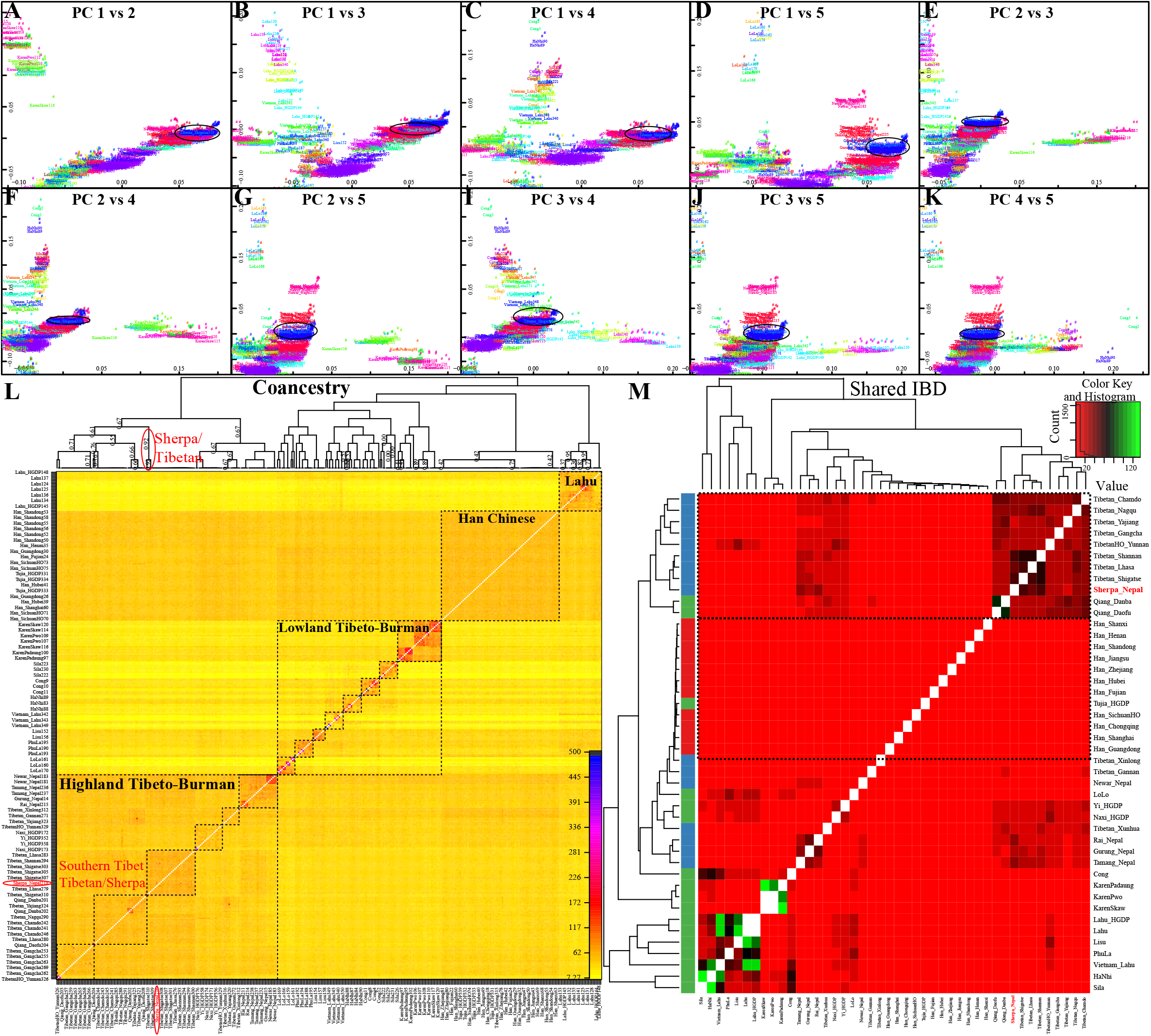
Shared ancestry among 44 Sino-Tibetan-speaking populations based on shared ancestry chunks. (A-I) IBD-based genetic relationships based on the top five components. (K) Coancestry inferred from the FineStructure-reconstructed IBD numbers. (L) IBD fragments among 44 populations inferred from the fineIBD analysis.

The patterns of population structure among the 18 populations revealed from FineStructure-based PCA showed a clear population substructure consistent with their ethnic or geographical divisions. PCA plots from the two components showed five genetic homogeneous clusters, including Newar, Taman and Gurung, Sherpa and Rai, Yi, and the Han Chinese (**Figure 7A**). the third component separated the Rai and Sherpa, and the fourth component separated northern and southern Han Chinese populations. Generally, Sherpas possessed closer relationships with Tamang, Rai and Gurung compared with others (**Figure 7B-C**). These genetic affinities between Sherpa and Nepal Rai and Tamang were further confirmed via the heatmap and clustering patterns of pairwise coincidence (**Figure 7D**). Nepal populations were clustered into genetically homogeneous populations consistent with their ethnic divisions, and Han Chinese were divided into northern, central and southern groups. Lowland Yi was clustered separately (**Figure 7D**). Generally, clustering patterns based on the coancestry matrix in ChromoPainter and FineStructure analyses revealed finer-scale population substructures among involved Sino-Tibetan-speaking populations and indicated that the target Nepal Sherpa had the closest genetic relatedness with Tibetan_Shigatse.

**Figure 7.**
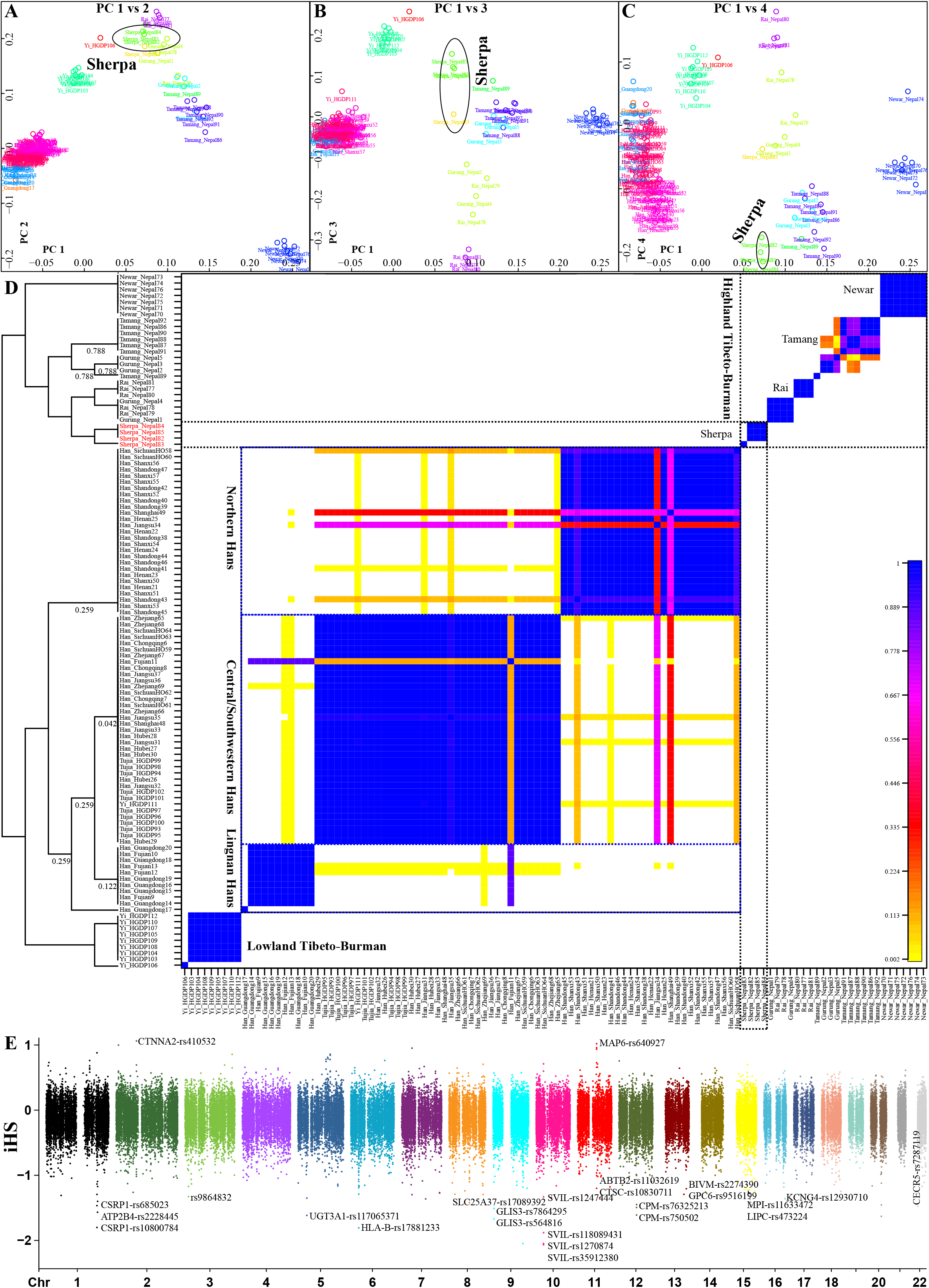
Fine-scale population structure among 18 Sino-Tibetan-speaking populations based on the reconstructed shared IBD numbers. (A-C) Patterns of genetic relationships based on the top five components. (D) Pairwise coincidence among 18 populations inferred from the fine-structure (fs) analysis. (E) Natural selection signatures among Sherpas and their neighbors from Nepal.

We also explored the potential natural-selection signals in the Nepal Sherpa based on the phased haplotypes using the integrated haplotype score (iHS), nSL, and integrated haplotype homozygosity pooled (iHH12). Positive selection signals were observed on chromosomes 1, 5, 6, 9, 10, 12, and 22 (**Figure 7E**), these observed SNPs were correlated with Cysteine and Glycine-Rich Protein 1 (CSRP1), UDP Glycosyltransferase Family 3 Member A1 (UGT3A1), Major Histocompatibility Complex, Class I, B (HLA-B), GLIS Family Zinc Finger 3 (GLIS3), Supervillin (SVIL), Carboxypeptidase M (CPM), and Cat Eye Syndrome Chromosome Region, Candidate 5 (CECR5), respectively. However, we did not observe variants that were in association with adaptive traits in the Nepal Sherpa, a further whole-genome sequencing project based on the larger sample size and single-molecule real-time sequencing technology should be conducted focused on Sherpa people in the next step, as the genetic findings in Tibetans [49].

## 4. Discussion

The Tibetan Plateau is widely known as the Third Pole of the world and is characterized by extreme climatic and environmental conditions [50]. As one of the most challenging environments ever occupied by anatomically modern humans (AMHs) [51], the timing and mechanisms of the colonization of Tibetan highlanders (Tibetans, Sherpas, Monpas, Lhobas, and Dengs) have attracted wide attention from various research fields, including genetics, palaeogenomics, archeology, anthropology, and geology [3, 9, 52–55]. Over the last few decades, with the deepening of researches in these fields, remarkable progress has been made in the exploration of the demographic history of Tibetans [52, 53, 55, 56]. However, as one of the high-altitude adaptive populations living around the Himalayas, the demographic history and fine-scale genetic background of Sherpas need to be comprehensively explored. The previous genetic analyses focused on the peopling of Tibetan Plateau mainly based on allele-based sharing ancestry or only modern genomes without the comprehensively comparative analysis based on both modern and ancient references [9, 57, 58]. Our recent population genetic analyses have started to directly explore the direct genetic contribution from ancient East Asians to modern Tibetan Plateau Tibetans based on ancient genomes, but the genetic history of Sherpa people and their forensic characteristics still need to be further studied [59–61]. In the present study, we first performed an InDel-based forensic and population genetic study to explore the genetic relatedness between Sherpa highlanders and worldwide reference populations. Furthermore, we conducted genome-wide SNP-based analyses among 382 modern and ancient populations to dissect the fine-scale genetic structure of Sherpa people based on the shared patterns of independent SNPs or reconstructed haplotypes.

The results of tests of HWE revealed that two InDel loci showed a departure from HWE after applying Bonferroni correction, which may be attributed to the population substructure, purifying selection, inbreeding, or copy number variation [62], consisting with high IBD fragments within Sherpa people. The allele frequency distributions and corresponding forensic parameters of 57 A-InDels in the studied Dingjie Sherpa indicated that several InDels (such as rs10607699, rs145577149, and rs66477007, among others) were not polymorphic in the target Dingjie Sherpa, but the values of TDP and CPE demonstrated that the AGCU InDel 60 kit is suitable for forensic individual identification in the Dingjie Sherpa. The InDel-based close genetic affinity between the Dingjie Sherpa and high-altitude Tibeto-Burman-speaking groups (especially Tibetans) supported that they originated from the same ancestral population or substantial gene flow among Tibetan highlanders had occurred [4, 5, 9]. The ancestral composition obtained from model-based ADMIXTURE analysis revealed that the European-dominant and Li-dominant ancestries might be absent in the gene pool of the Dingjie Sherpa, indicating that its ancestral components were mainly derived from East Asian-related ancestry. Additionally, we did not observe gene flow events from proxy sources into the Dingjie Sherpa, which suggested that more InDel-based genetic profiles from geographically/ethnically/linguistically different populations and genome-wide data from worldwide modern and ancient populations are indispensable to explore the fine-scale genetic makeup of the Sherpas.

The patterns of genetic clustering inferred from the genome-wide SNP-based PCA, ADMIXTURE, and TreeMix analyses revealed the population substructures among Sino-Tibetan-speaking groups in East Asia, Southeast Asia, and South Asia, and also indicated that the target Sherpas had close genetic affinities with high-altitude Tibetans and Nepal ancient individuals but had relatively distant genetic relationships with modern Nepalese populations, which supported the genetic findings that geographically different Tibeto-Burman-speaking groups have experienced complex and extensive admixtures [7, 8, 53] and the Sherpas possessed limited gene flow from surrounding Nepalese populations [7]. Furthermore, these genetic observations highlighted the long-term stability of the Tibetan highlanders’ genetic makeup [33]. Genetic similarity and continuity between the Sherpas and ancient populations and Tibetan groups on the Tibetan Plateau were further confirmed by their excess shared alleles revealed in *f*-statistics, qpWave/qpAdm-based admixture models, and qpGraph-based evolutionary topology. Moreover, we found that Lajia_LN-related ancestry contributed the most to the gene pool of modern Sherpa and Tibetan populations, suggesting that Neolithic millet farmers played a pivotal role in the formation of modern Sherpas and Tibetans, which was consistent with previous genetic findings [36, 53, 59]. The haplotype-based genetic observations further confirmed the recent common ancestor sharing between the Sherpas and Tibetans. Additionally, we observed several positive selection signals not associated with high-altitude adaptation across the genomes of Sherpas, which may result from the limited sample sizes.

## 5.Conclusion

Sherpa people are one of the officially unrecognized ethnolinguistic groups, like the Gejia, Dongjia, and Xijia in Guizhou province. Here, we provided genetic evidence from both InDels and genome-wide SNPs and suggested that Sherpa people shared a common admixture and evolutionary history with core Tibetans and Tibeto-Burman-speaking populations from surrounding regions. We identified the genetic affinity between Sherpas and modern Tibetans and ancient populations on the Tibetan Plateau, and this genetic homogeneity among them was further confirmed in the PCA, ADMIXTURE, Fst and allele-sharing profiles in *f*-statistics and haplotype-based shared ancestry fragments. Consistent with their modern and ancient neighbors (such as Chokhopani and Tibetans), Sherpa people possessed strong lowland East Asian affinity and was formed via the admixture of major ancestry related to the Yellow River Basin Neolithic millet farmers and minor from the indigenous ancestry related to the early East Asians.

## Acknowledgments

This study was supported by grants from the National Natural Science Foundation of China (31801040), Nanqiang Outstanding Young Talents Program of Xiamen University (X2123302), and Fundamental Research Funds for the Central Universities (ZK1144). The funders had no role in study design, data collection, and analysis, preparation of the manuscript, or decision to submit the manuscript for publication. GLH was supported by Project funded by China Postdoctoral Science Foundation (2021M691879). We thank all volunteers who participated in this project and also thank Wibhu Kutanan (Department of Biology, Faculty of Science, Khon Kaen University), Dang Liu and Mark Stoneking (Department of Evolutionary Genetics, Max Planck Institute for Evolutionary Anthropology) for sharing genotypes of modern populations from Vietnam, Thailand and Laos.

## Legends of Supplementary materials

**Figure S1.** PCA results based on genotype (**A~B**) and allele frequency distribution (**C~D**).

**Figure S2.** TreeMix analysis among Human Origin dataset.

**Figure S3.** Natural selection signals in Nepal populations.

